# Distinct Spatial Programs of Response versus Resistance in Non-Small Cell Lung Cancer after Neoadjuvant Chemoimmunotherapy

**DOI:** 10.64898/2026.04.05.716543

**Authors:** Seo Hye Park, Jaemoon Koh, Sungwoo Bae, Hongyoon Choi, Taeyoung Yun, Jihyeon Park, Bubse Na, Samina Park, Hyun Joo Lee, In Kyu Park, Chang Hyun Kang, Young Tae Kim, Kwon Joong Na

## Abstract

**Background:** Neoadjuvant chemoimmunotherapy (nCIT) has become a standard treatment for locally advanced resectable non-small cell lung cancer (NSCLC), yet the spatial biology underlying treatment resistance remains poorly understood. We used spatial transcriptomics to define the microenvironmental architecture of residual cancers in patients who did not achieve major pathologic response (non-MPR) compared with those who did (MPR).

**Methods:** Spatial transcriptomics was performed on 10 formalin-fixed paraffin-embedded (FFPE) tumor blocks (5 MPR, 5 non-MPR) obtained from 8 patients treated with nCIT. A deep learning algorithm was applied to detect viable residual cancer spots from treatment-induced fibrosis and necrosis. Spatial deconvolution, distance modeling, ligand-receptor analysis, and functional pathway scoring were integrated to characterize niche-specific programs.

**Results:** MPR cancer core displayed an immune-permissive remodeling environment with deep infiltration of cytotoxic CD8+ T cells, mature dendritic cells (LAMP3+, CCR7+), and active efferocytosis signaling (APOE–TREM2), alongside robust MHC class II expression. Non-MPR cancer core, by contrast, exhibited spatial immune exclusion: a dense fibroblast barrier reinforced by TIMP1–CD63 signaling and Treg-enriched boundaries physically restricted effector T cell access to the cancer core. Residual cancer cells in non-MPR samples maintained active cell cycling and independently upregulated cytochrome P450-mediated drug detoxification and DNA damage response pathways without inducing MHC class II expression — effectively decoupling intrinsic survival from immune recognition. The non-MPR core also showed a hyper-metabolic profile, including elevated glutathione metabolism consistent with antioxidant buffering against chemotherapy-induced oxidative stress.

TROP2 was broadly expressed across the non-MPR cancer core and co-localized with DNA damage response and nuclear factor erythroid 2-related factor 2 resistance signatures.

**Conclusions:** Residual cancer cores in non-MPR tumors appear to represent evolved resistant niches sustained by structural immune exclusion, metabolic rewiring, and DNA repair proficiency. These findings highlight the spatial co-localization of epithelial anchors, such as TROP2, with intrinsic resistance pathways, providing a structural rationale for developing novel precision therapeutic strategies to bypass stromal barriers and overcome the cancer core’s intrinsic repair capacity.

## Background

The combination of immune checkpoint inhibitors (ICIs) with platinum-based chemotherapy has established chemoimmunotherapy (CIT) as the standard of care for metastatic non-small cell lung cancer (NSCLC), delivering consistent survival benefits across multiple trials. [1] This success has extended to earlier disease stages: neoadjuvant CIT (nCIT) now improves event-free and overall survival in resectable NSCLC compared with chemotherapy alone. [2,3] Despite these advances, a substantial fraction of patients does not respond. The biological basis of this resistance — specifically, how residual cancer cells withstand the combined selective pressure of cytotoxic chemotherapy and immune reactivation — remains incompletely understood. [4,5] Identifying the vulnerabilities of these treatment-refractory clones is essential for developing rational salvage strategies.

The neoadjuvant setting offers a unique opportunity to study resistance biology. Unlike palliative treatment, where post-progression tissue is rarely available, surgical resection after nCIT provides the entire treated tumor bed for analysis. The International Association for the Study of Lung Cancer (IASLC) defines major pathologic response (MPR) as ≤10% viable residual tumor, a threshold validated as a surrogate for long-term survival. [6,7] The viable cancer cells persisting in non-MPR specimens (>10% residual cancer), therefore, represent resistant clones that have survived both cytotoxic and immune-mediated killing in vivo. Characterizing these cells and their surrounding microenvironment offers a direct route to understanding treatment failure.

Doing so, however, poses significant analytical challenges. After nCIT, the tumor bed is a heterogeneous mixture of necrotic debris, fibrotic regression beds, and variable immune infiltrates surrounding scattered cancer cores. Bulk transcriptomics dilutes the signal from rare viable cells with overwhelming fibrotic and necrotic noise. Single-cell RNA sequencing provides cellular resolution but sacrifices the tissue architecture needed to evaluate structural barriers to immune access. Spatial transcriptomics (ST) overcomes both limitations: by preserving spatial coordinates, ST enables mapping of surviving cancer cells in their native context and can distinguish whether treatment failure arises from physical immune exclusion by the stroma region, from cancer-intrinsic survival adaptations within a protected core, or from both.

In this study, we applied ST to compare the microenvironmental landscapes of MPR and non-MPR NSCLC following nCIT. To address the morphological ambiguity of treated tissues, we combined pathologist-guided macro-dissection with Cancer-Finder[8], a domain-generalization-based deep learning model, to computationally delineate the cancer-specific niche. Unlike morphology-based methods, Cancer-Finder’s ability to generalize across diverse transcriptomic contexts allowed for the robust identification of malignant spots even within the complex, therapy-induced fibrotic and necrotic areas of nCIT-treated. We hypothesized that resistance to nCIT is governed by highly organized spatial niches, driven by a combination of stromal-mediated immune exclusion and cancer-intrinsic survival programs. Furthermore, we postulated that if resistant cancer cells are physically shielded from immune infiltration, delivering cytotoxic payloads directly to the epithelial surface via antibody-drug conjugates (ADCs) could represent a rational precision strategy to bypass the stroma. To explore this, we mapped the spatial expression of TROP2 (TACSTD2)—a leading epithelial target for emerging ADCs in NSCLC [9]—alongside intrinsic resistance signatures. Our goal was to define the spatial architecture of nCIT resistance and to evaluate how mapping epithelial surface targets can inform the development of novel salvage strategies to dismantle evolved resistant niches.

## Materials and Methods

### Study Ethics and Patient Cohort

This study was approved by the Institutional Review Board of Seoul National University Hospital (IRB approval number: H-2009-081-1158, approval date: Sep 18th, 2020) and conducted in accordance with the Declaration of Helsinki.

We enrolled eight patients diagnosed with non-small cell lung cancer (NSCLC) who received nCIT between October 2022 and December 2023. All patients received a combination of platinum-based chemotherapy and PD-1/PD-L1 inhibitors, followed by surgical resection. Pathological response was evaluated by a board-certified thoracic oncology pathologist (J.K.) according to the IASLC recommendations [6]. MPR was defined as ≤10% viable residual cancer, while non-MPR was defined as >10% viable residual cancer. Pathological complete response cases were excluded as they lack residual cancer cells to reflect resistance mechanisms. A total of 10 representative formalin-fixed paraffin-embedded (FFPE) cancer blocks (5 MPR, 5 non-MPR) were selected for ST analysis.

### ST Data Generation and Quality Control

ST data generation was performed using the Visium CytAssist Spatial Gene Expression Kit (10x Genomics, Pleasanton, CA, USA). To evaluate the RNA quality of the FFPE samples, we assessed the percentage of RNA fragments greater than 200 nucleotides (DV200) according to the manufacturer’s protocol. Only samples with a DV200 value exceeding 30% were included in the study to ensure sufficient transcriptomic data quality. FFPE blocks were sectioned at a thickness of 5 µm and placed on positively charged glass slides. After deparaffinization, H&E staining was performed, and the slides were digitally scanned. For precise spatial analysis, a board-certified thoracic oncology pathologist (J.K.) manually selected the optimal capture areas (6.5 x 6.5 mm), specifically targeting regions with high viable cancer cellularity while strictly avoiding areas of extensive necrosis or non-specific tissue debris. The sections were then processed for RNA probe hybridization using the Visium Human Transcriptome Probe Kit v2. Probe pairs were ligated and transferred to a Visium CytAssist slide with a 6.5 x 6.5 mm capture area. Libraries were constructed following the manufacturer’s instructions and sequenced with a target depth of 50,000 read pairs per spot.

### Pre-Processing and Data Integration

Raw sequencing data were processed using SpaceRanger software to align reads to the human reference genome (GRCh38). Downstream analysis was performed with Python (v3.10). To integrate 10 ST datasets and mitigate batch effects, we employed single-cell Variational Interference [10]. The raw counts were normalized to a target sum of 1×10^4^ and log-transformed using the log1p function. Highly variable genes were identified using the Seurat v3 flavor (n=3,000) based on the normalized expression layer.

### Transcriptome profile-based cancer region definition

To precisely define the viable cancer regions within the selected ST capture areas, we had to overcome the inherent limitations of conventional morphological annotation. Because residual cancer cells post-nCIT are frequently fragmented and obscured by extensive therapy-induced fibrosis and necrosis, visual assessment alone is insufficient for capturing the precise boundaries of isolated cancer nests. To resolve this, we utilized Cancer-Finder, a deep learning-based model, to computationally delineate the cancer-specific niche at a near-single-spot resolution. By classifying spots based on their global transcriptomic profiles rather than morphology, this approach allowed us to effectively isolate the true transcriptomic signals of viable cancer regions from transcriptionally inactive debris or purely fibrotic areas.

### Spatial Niche Modeling and Distance Analysis

To estimate the abundance of specific cell types within each Visium spot, we performed spatial deconvolution using cell2location [11]. As a reference single-cell RNA sequencing dataset, we utilized the comprehensive NSCLC data (GEO accession number: GSE131907) [12]. Specifically, the annotation of immune and stromal cell subtypes was informed by the high-resolution phenotypic atlas provided by Lambrechts et al. [13], ensuring precise spatial deconvolution of the lung tumor microenvironment. To systematically characterize the spatial architecture of the tumor microenvironment (TME), we categorized the tissue into two primary compartments: the cancer region (viable cancer spots identified by Cancer-Finder) and the stroma region (all remaining non-cancer spots).

Spatial distance was calculated as the Euclidean distance from each spot to the nearest cancer boundary defined by Cancer-Finder using Shapely (v2.1.1) and GeoPandas (v1.1.1), then normalized to a percentage scale (0% at boundary, 100% at innermost core) for cross-sample comparison. This integrated spatial framework served as the basis for all subsequent niche-specific differential expression and Generalized Estimating Equations (GEE) analysis.

### Functional Signature and Pathway Analysis

Functional activity was assessed using gene set variation analysis or add-on module scores for specific signatures encompassing immune function (cytotoxicity, exhaustion, regulatory T cell (Treg), and antigen presentation), resistance axes (DNA damage response including homologous recombination (HR), non-homologous end joining (NHEJ), and the nuclear factor erythroid 2-related factor 2 (NRF2) pathway), and metabolism (glycolysis, oxidative phosphorylation (OXPHOS), tricarboxylic acid (TCA) cycle, and glutathione (GSH) metabolism)

Differentially expressed genes (DEGs) were identified via Wilcoxon rank-sum tests with Benjamini-Hochberg correction. Over-representation analysis was performed using Gseapy (v1.1.9) against Reactome 2022 and KEGG 2021 human databases. Pathway overlaps were visualized as networks using NetworkX (threshold ≥ 5 gene overlap). Decoupler (v2.1.1) was utilized for auxiliary enrichment calculations.

### Ligand-Receptor Interaction and Spatial Pattern Analysis

Spatially-resolved ligand-receptor interactions were analyzed using the LIANA package (v1.6.0). [14] We utilized the rank_aggregate method to identify dominant communication axes and specifically modeled stroma-to-cancer cross-talk. Global spatial autocorrelation was evaluated using Moran’s I via Squidpy (v1.6.5). To pinpoint resistance hotspots and quantify their spatial co-localization with TROP2 (TACSTD2), we utilized quantile-based thresholding and spatial embedding maps.

### Statistical Analysis

To account for the intra-patient correlation of spots, we employed GEE method. We used a Gaussian family with an exchangeable correlation structure, treating the pathological response group (MPR vs. non-MPR) as the primary fixed effect. GEE analysis was performed separately for cancer and stroma regions to identify niche-specific adaptations. All p-values were adjusted for multiple testing.

## Results

### Cohort Characteristics and Precise Spatial Mapping of residual Tumor Niches

The clinical characteristics of the study cohort are detailed in Table 1. H&E-stained images revealed a highly heterogeneous microenvironment where treatment-induced fibrosis, necrosis, and inflammatory infiltrates were intricately mixed (Supplementary Figure 1A). To resolve this morphological ambiguity, the application of the Cancer-Finder algorithm provided a sharp qualitative contrast between the two groups based on their transcriptomic profiles (Supplementary Figure 1B). In the MPR group, reflecting a successful therapeutic response, Cancer-Finder-defined cancer regions were identified only as small, sparse, and scattered spots isolated within vast fibrotic areas. In contrast, non-MPR specimens were characterized by large, expansive, and densely clustered islands of cancer regions.

**Table 1.** Clinicopathological characteristics of the sample cohort.

| **Patient ID** | **Group** | **Histology** | **Clinical Stage (Pre-nCIT)** | **Pathologic Stage (Post-nCIT)** | **PD-L1 (TPS)** | **SUVmax (Pre)** | **SUVmax (Post)** |
| --- | --- | --- | --- | --- | --- | --- | --- |
| **P-01, P-02** | **MPR** | SqCC | T2bN0M0 | T1aN0M0 | 0-1% | 17.6 | 2.9 |
| **P-03** | **MPR** | SqCC | T4N0M0 | T0N0M0 | 2% | 19.5 | 5.4 |
| **P-04, P-05** | **MPR** | Adeno | T3N1M0 | T1aN0M0 | 60-80% | 17.7 | 4.3 |
| **P-06** | **non-MPR** | Adeno | T4N1M0 | T3N1M0 | 5% | 15.2 | 19.1 |
| **P-07** | **non-MPR** | SqCC | T3N1M0 | T1bN0M0 | 0% | - | - |
| **P-08** | **non-MPR** | Adeno | T2aN0M0 | T2bN0M0 | 3% | - | - |
| **P-09** | **non-MPR** | SqCC | T4N1M0 | T1bN1M0 | 90% | 16.4 | 7.6 |
| **P-10** | **non-MPR** | SqCC | T3N0M0 | T2aN1M0 | 30% | 11.7 | 16.5 |

*Adeno, adenocarcinoma; nCIT, neoadjuvant chemoimmunotherapy; PD-L1, programmed death-ligand 1; SqCC, squamous cell carcinoma; SUVmax, maximum standardized uptake value; TPS, tumor proportion score.*

To characterize the molecular drivers of these distinct architectures, we identified the top differentially expressed genes (DEGs) between MPR and non-MPR groups in both the cancer and stroma regions. In MPR samples, the cancer regions were characterized by the upregulation of genes involved in antigen presentation and immune modulation (e.g., CD74, B2M) and macrophage-associated lipid metabolism (APOE, SPP1). The corresponding stroma region in MPR showed a robust infiltration of plasma cells, evidenced by high expression of immunoglobulin genes (IGKC, IGHG1) and markers of secretory activity (MZB1, JCHAIN), alongside active extracellular matrix (ECM) remodeling (COL1A1, SPARC). Conversely, non-MPR cancer regions were dominated by mitochondrial stress-related transcripts (e.g., MT-CO3, MT-ATP6), while their surrounding stroma regions retained lung-specific markers (SFTPC, SCGB3A1), reflecting a less remodeled and potentially immunosuppressive microenvironment. These baseline transcriptomic divergences establish the core molecular identities of the responding and resistant niches, serving as the direct foundation for our subsequent spatial interaction and distance-based modeling (**Supplementary Table 1**).

### Non-MPR tumors exhibit spatial immune exclusion enforced by a dense stromal barrier

To characterize the spatial distribution of tumor microenvironment (TME), we first estimated the proportions of major cell types across the cohort (**Figure 2A**). In the MPR group, immune cell populations, including myeloid cells, B cells, and T lymphocytes were significantly more abundant within the cancer region compared to the non-MPR group, which was conversely dominated by epithelial cells in the cancer region. While global proportions provided an overview, stratified analysis of the cancer and stroma regions revealed distinct spatial partitioning (**Figure 2B**).

**Figure 1.**
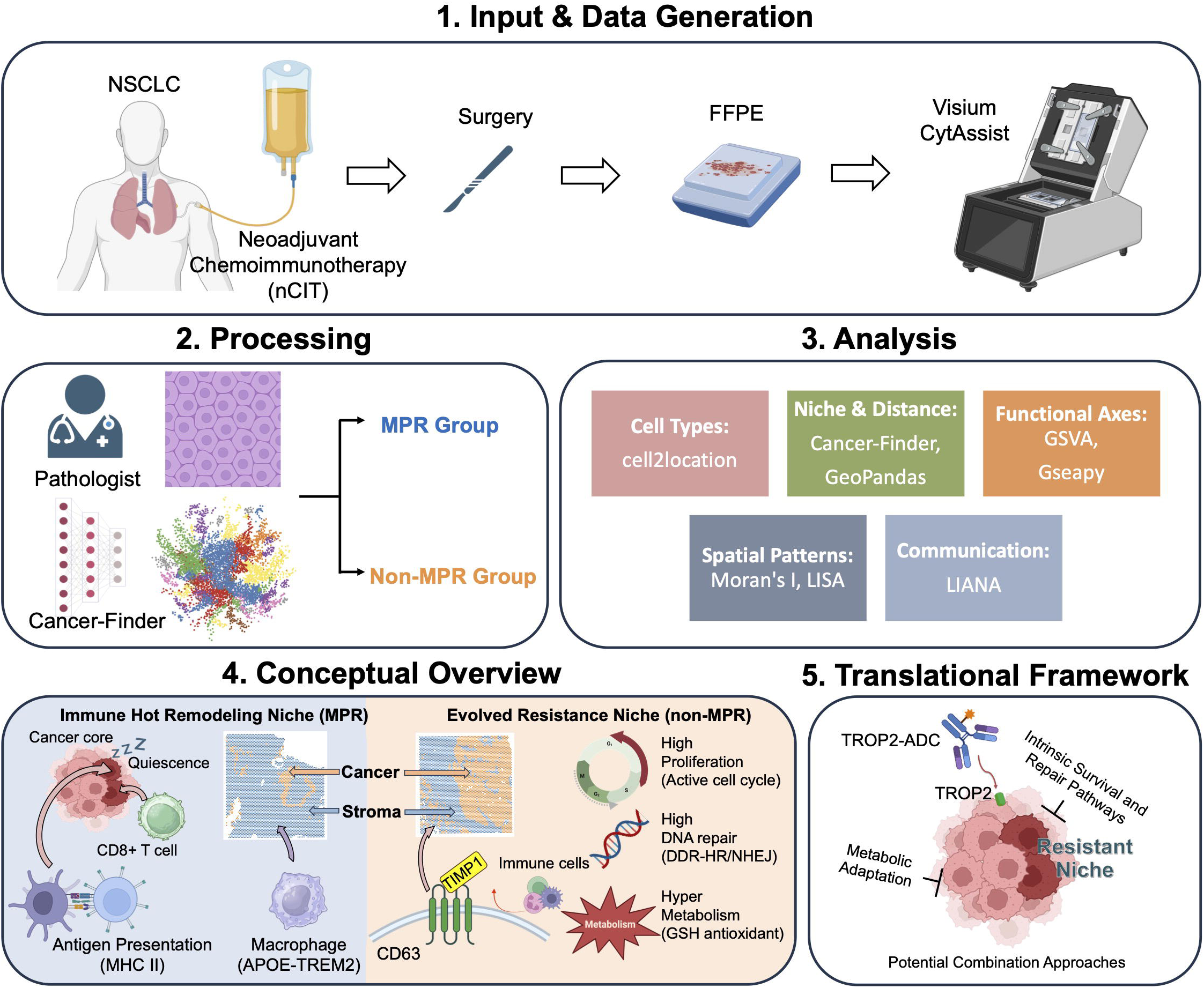
Study workflow and spatial landscape of therapy-induced residual niches. (1) Input and data generation: Formalin-fixed paraffin-embedded (FFPE) tumor blocks were obtained from non-small cell lung cancer (NSCLC) patients who underwent surgical resection after neoadjuvant chemoimmunotherapy (nCIT). Spatial transcriptomic data were generated using the Visium CytAssist platform (10x Genomics). (2) Processing: To address the morphological complexity of treated tissues, pathologist-guided macro-dissection was combined with the transcriptome-based Cancer-Finder algorithm to define viable residual cancer regions within treatment-associated fibrosis and necrosis. (3) Analysis: Integrated spatial analyses included cell-type deconvolution, boundary-based distance modeling, pathway scoring, ligand–receptor inference, and spatial statistics. (4) Conceptual overview: Schematic comparison of the spatial programs observed in MPR and non-MPR tumors. MPR lesions were associated with immune infiltration, antigen-presentation features, and quiescent residual cancer cells, whereas non-MPR lesions showed fibroblast-enriched boundary features and cancer-core programs associated with proliferation, DNA repair, and metabolic activity. (5) Therapeutic rationale: TROP2 retained in the resistant epithelial compartment is depicted as a candidate spatial anchor for antibody-drug conjugate delivery to residual disease. The spatial overlap between TROP2 and intrinsic resistance-related programs provides a rationale for future evaluation of combination approaches in residual non-MPR disease.

**Figure 2.**
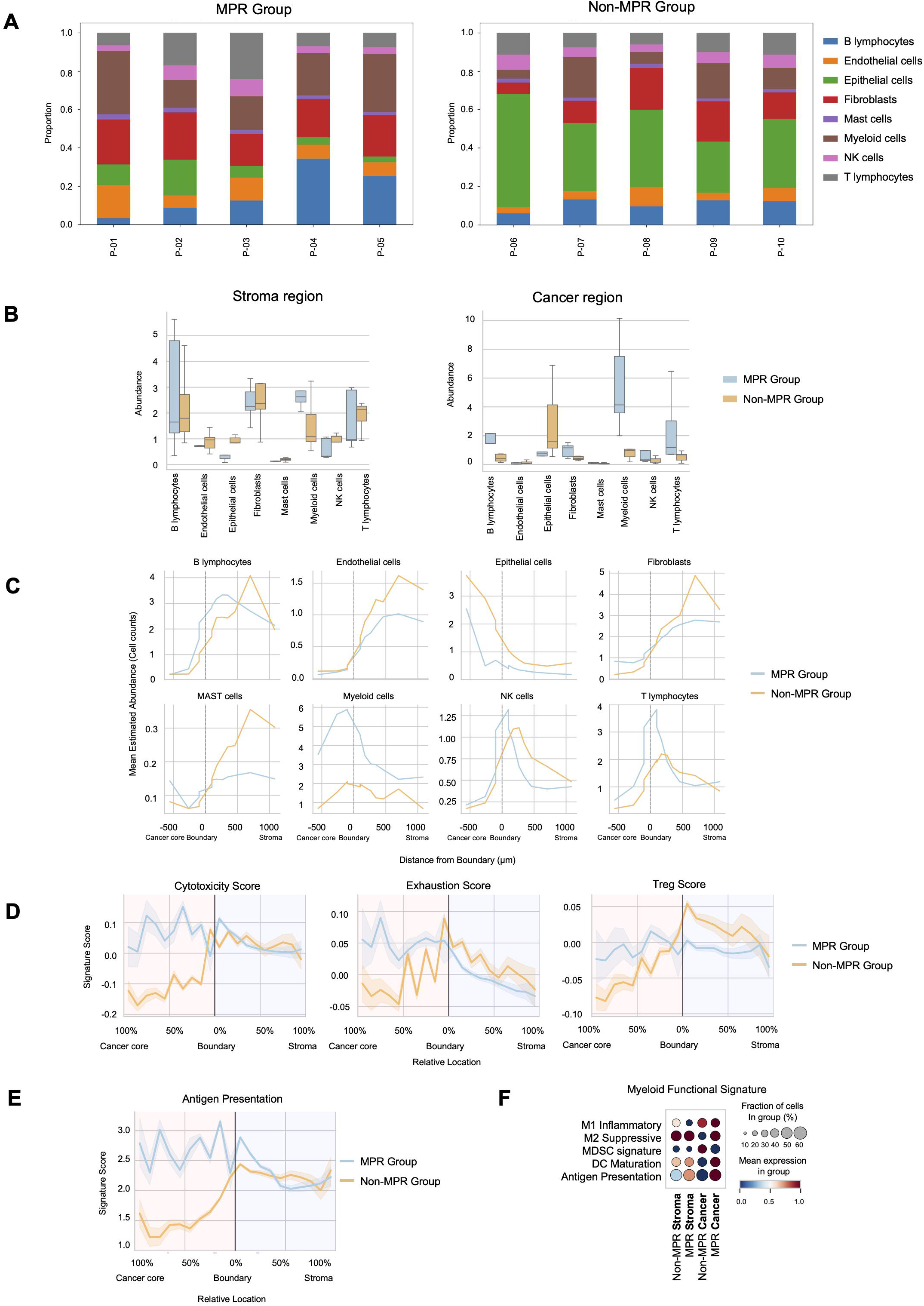
Spatial architecture of immune infiltration and immune exclusion in MPR and non-MPR tumors. (A) Stacked bar plots show the estimated proportions of major epithelial, immune, and stromal cell types in each sample. (B) Box plots compare cell-type abundance between stroma regions and Cancer-Finder-defined cancer regions according to response group. (C) Distance-based profiles show mean cell-type abundance as a function of distance from the tumor boundary (0 μm), with negative values representing the cancer-core side and positive values representing the stroma region side. (D-E) Boundary-aligned trends of cytotoxicity, exhaustion, Treg, and antigen-presentation signatures across the cancer-to-stroma axis. (F) Dot plots summarize myeloid functional signatures across group-by-region compartments, with dot size indicating the fraction of spots and color indicating mean expression. Collectively, these analyses depict a more immune-permissive architecture in MPR and a more immune-excluded configuration in non-MPR tumors.

To further delineate the structural differences, we performed a distance-based density analysis from the tumor boundary into the core and out toward the stroma region (ranging from -500 μ to 1000 μ) (**Figure 2C**). In both MPR and non-MPR groups, the density of immune cells (myeloid cells, B cells, and T lymphocytes) exhibited a decreasing gradient from the stroma region into the cancer core. However, a critical difference was observed in the retention of these cells: the MPR group maintained a significantly higher and more sustained presence of immune infiltrates deep within the cancer core, whereas these cells were nearly absent in the non-MPR core. (**Figure 2C**). Regarding the stromal architecture, although fibroblasts were present in the cancer cores of both groups, the non-MPR group exhibited a sharp surge in fibroblast density specifically within the stroma and at the tumor boundary, These findings suggests that in non-MPR tumors, fibroblasts form a dense physical barrier that restricts the influx of immune cells into the epithelial-dense cancer core, reinforcing an ‘immune-excluded’ phenotype. In the non-MPR cancer region, T cells were found to be extremely sparse, with their numbers increasing only as the distance from the boundary toward the stroma region increased. This consistent pattern of exclusion was observed across all lymphoid subtypes, including cytotoxic CD8+ T cells, exhausted CD8+ T cells, and NK cells. In contrast, these cytotoxic and functional effector cells were found to be well-distributed throughout the cancer core in MPR patients (**Supplementary Figure 2**).

We then evaluated the functional characteristics of the immune TME by calculating signature scores for cytolytic activity, exhaustion and Tregs based on their distance from the tumor boundary (**Figure 2D**). In the MPR group, the cancer region was characterized by concurrently high levels of cytotoxicity and exhaustion scores. This suggests that immune cells did not merely remain in the stroma region but successfully infiltrated the cancer core to engage the cancer cells. Such a profile likely reflects a state of chronic activation, where prolonged immune engagement may have ultimately led to functional exhaustion. Conversely, the non-MPR group displayed elevated Treg scores concentrated around the boundary, which likely acts as an additional immunological barrier limiting the entry of effector T-cells (**Figure 2D and Supplementary Figure 3**). Lastly, we investigated the capacity for antigen recognition within these niches. The MPR cancer core was significantly enriched with mature dendritic cells (DC) (LAMP3- and CCR7-expressing DCs) and high antigen presentation signatures, including MHC class II (**Figure 2E, 2F, and Supplementary Figure 2 and 3**). The co-localization of MHC class II expression with mature DCs suggests a TME that facilitates active antigen processing and presentation, thereby allowing T cells to effectively recognize cancer cells. In non-MPR cancer cores, the functional void of these antigen-presenting cells and associated MHC signatures further explains the failure of the immune system to dismantle the resistant epithelial-dense core.

**Figure 3.**
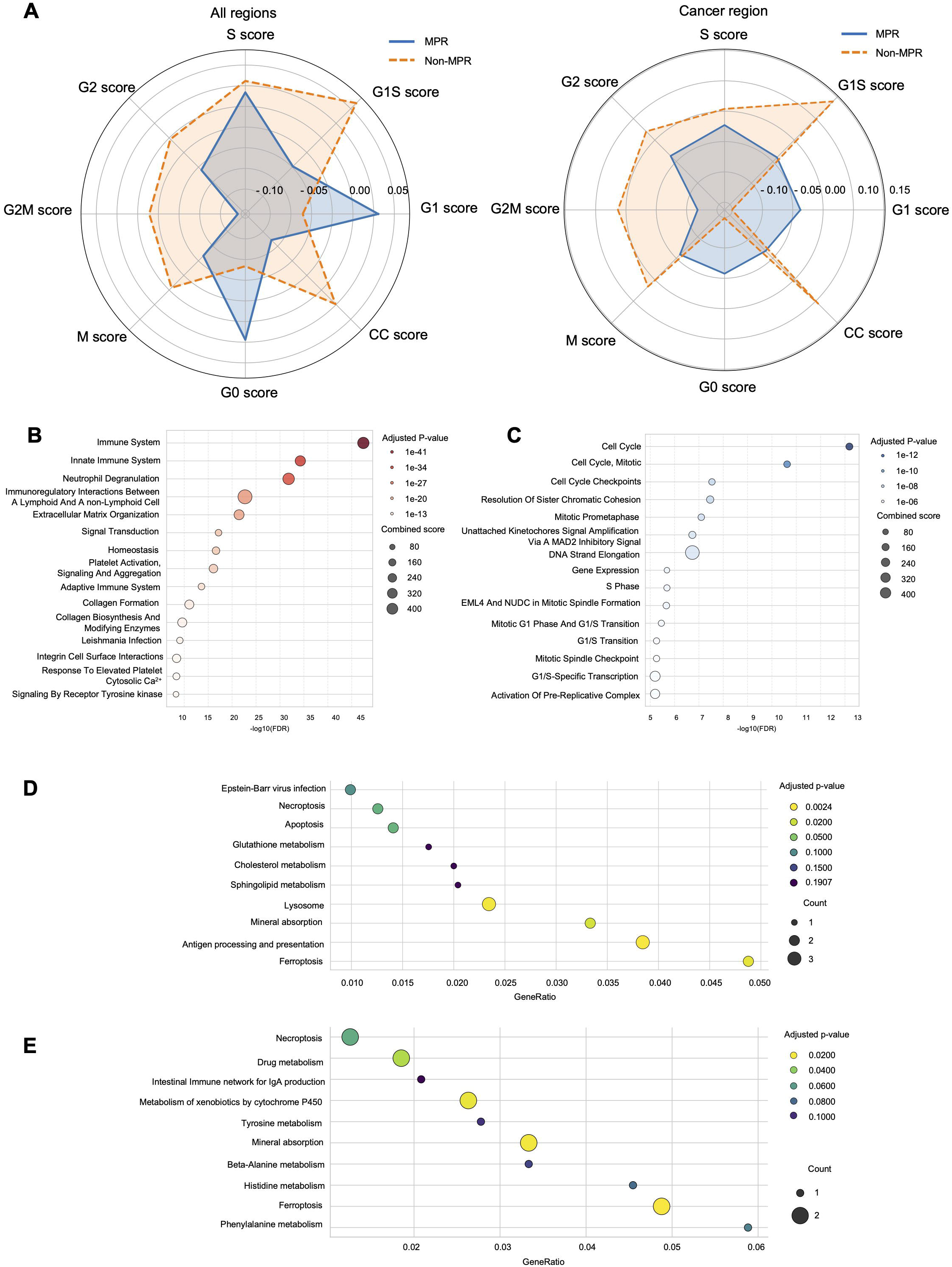
Cell-cycle states and pathway programs in residual cancer regions. (A) Radar plots compare cell-cycle phase scores across all regions and Cancer-Finder-defined cancer regions in MPR and non-MPR groups. (B-C) Bubble plots show the top enriched Reactome pathways in MPR (B) and non-MPR (C) cancer regions, with the x-axis representing -log10(FDR) and dot size representing the combined score. (D-E) KEGG pathway enrichment in MPR (D) and non-MPR (E) cancer regions, with the x-axis showing GeneRatio, dot size indicating gene count, and color indicating adjusted p value. In aggregate, MPR-associated cancer regions show features compatible with quiescence and tissue/immune remodeling, whereas non-MPR cancer regions retain proliferative, cell-cycle, and xenobiotic or metabolic programs.

### Non-MPR residual tumors exhibit active cell cycling and intrinsic detoxification

To interrogate the biological status of residual cancer cells surviving nCIT, we analyzed cell cycle programs based on RNA transcript levels specifically within the defined cancer regions of the MPR and non-MPR (**Figure 3A**). Radar plots of cell cycle phase scores revealed that residual cells in the MPR group were primarily in a quiescent, non-proliferative state, characterized by high G0/quiescence scores and significantly lower scores for DNA synthesis and mitosis phase. In contrast, the non-MPR group exhibited a shift toward active cell cycling, with markedly higher scores for the S and G2M phases and a corresponding reduction in G0 scores across the entire cancer region. These findings indicate that while MPR cancer cells remain in a state of dormancy or growth arrest, non-MPR cancer cells maintain an active proliferative state program, continuing to divide despite the selective pressure of nCIT.

To further delineate the functional landscape of these surviving clones, we performed Reactome and KEGG pathway enrichment analysis restricted to the cancer regions. In MPR cancer regions, these analyses were predominantly enriched for innate immune activation, antigen presentation, and tissue remodeling (e.g., ECM organization and programmed cell death). In stark contrast, non-MPR cancer regions were uniquely characterized by intrinsic survival programs, including active cell-cycle progression, RNA transcription, and xenobiotic/drug metabolism (Figure 3B–3E). The full statistical output for these Reactome and KEGG pathway enrichment analyses, including GeneRatios and adjusted p-values, is provided in **Supplementary Table 2**; stromal region pathway enrichment is shown in **Supplementary Figure 4**.

Notably, KEGG pathway analysis revealed markedly divergent functional programs between the two groups (**Figure 3D, 3E**). In the MPR cancer region, pathways associated with programmed cell death (apoptosis, necroptosis, ferroptosis) and immune recognition (antigen processing and presentation) were highly enriched, reflecting ongoing immunogenic cell clearance. Conversely, the non-MPR cancer regions were uniquely characterized by active detoxification and metabolic rewiring mechanisms, highlighted by ‘drug metabolism’ and ‘metabolism of xenobiotics by cytochrome P450’. This functional decoupling suggests that within the resistant niche in non-MPR, the functional focus has shifted toward drug resistance and intrinsic survival programs. By independently managing intracellular drug toxicity and oxidative stress without triggering MHC class II expression, the resistant cancer core secures a stealth survival advantage—maintaining cellular homeostasis against ongoing therapeutic pressure while remaining completely hidden from the excluded immune TME.

### Non-MPR cancer cores exhibit profound metabolic flexibility and heightened DDR

To investigate whether specific metabolic adaptations contribute to cancer resistance to nCIT, we analyzed metabolism signatures from the ST data. Metabolic pathway scores revealed that non-MPR tumors are in a hyper-metabolic state. It was consistent with the post-nCIT ^18^F-fluorodeoxyglucose positron emission tomography (PET) results, as non-MPR patients showed significantly higher maximum standardized uptake value, though they underwent full cycles of nCIT. The metabolic profile of non-MPR tumors was characterized by simultaneous increases in Glycolysis, OXPHOS, the TCA cycle, FAO, and GSH metabolism (**Figure 4A, 4B**). Elevated GSH metabolism may reflect antioxidant buffering to neutralize chemotherapy-induced oxidative stress, potentially supporting cell survival and maintaining viability under therapeutic pressure.

**Figure 4.**
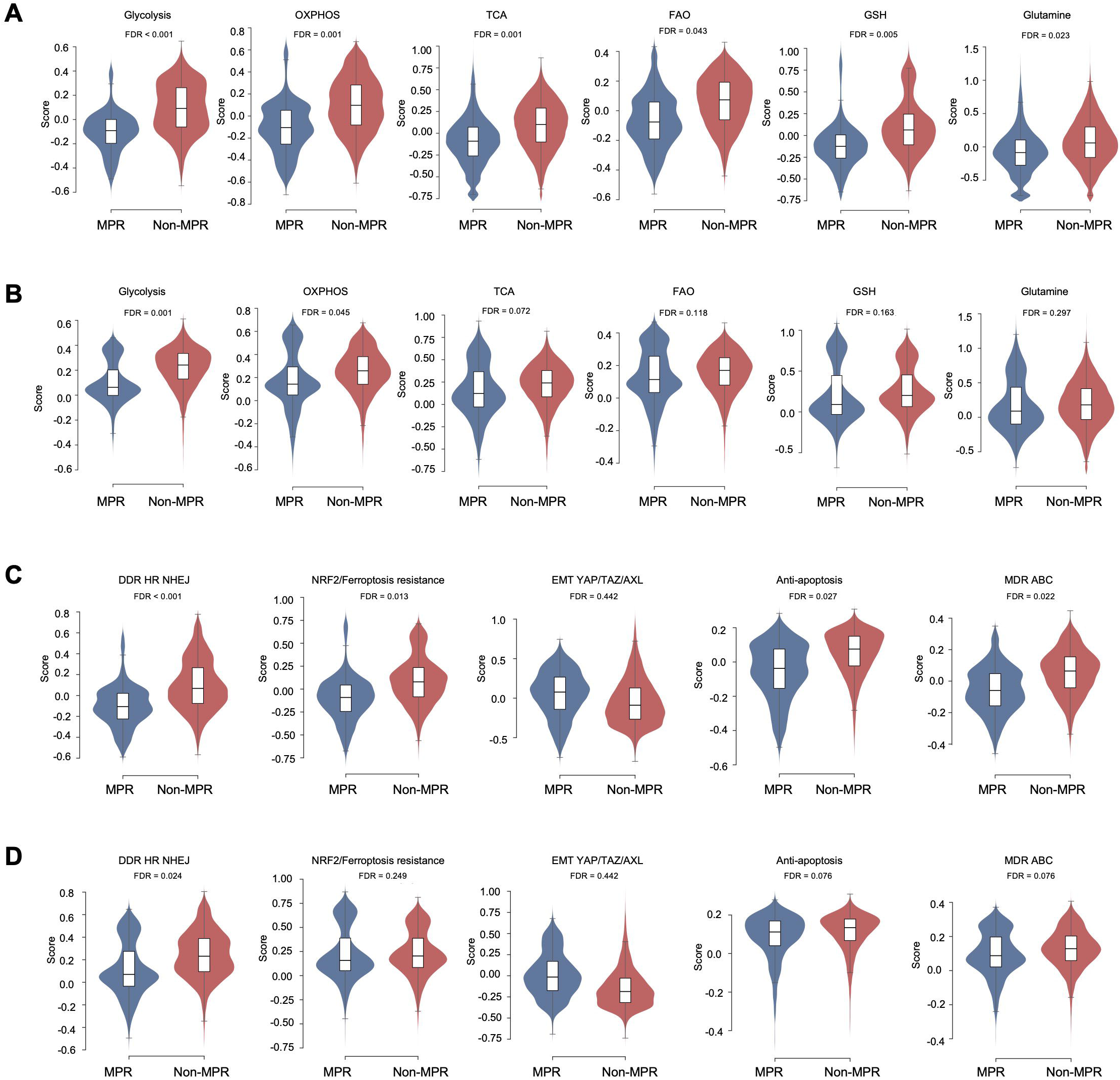
Metabolic and resistance-related programs in residual tumor regions. (A-B) Violin plots compare metabolic pathway scores across all spots (A) and Cancer-Finder-defined cancer regions (B), including glycolysis, oxidative phosphorylation, tricarboxylic acid cycle, fatty acid oxidation, glutathione metabolism, and glutamine metabolism. (C-D) Violin plots compare resistance-related signature scores across all spots (C) and cancer regions (D), including DNA damage response, NRF2/Ferroptosis resistance, EMT/YAPTAZ/AXL, anti-apoptosis, and MDR-ABC programs. FDR values are shown above each comparison. The separation between groups is more pronounced in cancer regions than in the overall tissue, supporting enrichment of cancer-intrinsic metabolic and repair programs in non-MPR residual disease.

Furthermore, non-MPR cancer regions showed significantly higher scores for DDR HR and NHEJ pathways compared to MPR (FDR < 0.05) (**Figure 4D**). This suggests that non-MPR cancer cells may possess DNA repair proficiency allowing them to effectively mitigate the double-strand breaks typically induced by platinum-based agents. In addition to DDR, we examined other resistance-related programs, including the NRF2/Ferroptosis resistance pathway, epithelial-mesenchymal transition (EMT), Anti-apoptosis signature, and Multi-Drug Resistance (MDR) involving ATP-Binding Cassette (ABC) transporters. While these modules also exhibited trends toward higher expression in the non-MPR cancer region, they did not reach statistical significance. Interestingly, when analyzed separately in cancer and stroma regions, these functional differences were notably attenuated in the stroma region. This observation supports the concept of the resistance niche, an emergent TME formed by the integration of active cancer-intrinsic survival programs and a protective stromal architecture (**Figure 4C, 4D**). Metabolic rewiring and chemo-resistance signatures were less significant when analyzed specifically within the stromal regions (**Supplementary Figure 5 and 6**).

### Spatial communication networks reveal active efferocytosis in MPR and fibrotic barrier maintenance in non-MPR tumors

To understand how these distinct spatial niches are actively maintained, we performed ligand-receptor interaction analysis using LIANA. In both groups, a CD44-centered signaling axis (e.g., VIM–CD44, FN1–CD44, COL1A1–CD44) was highly prominent across both cancer and stroma regions (**Figure 5A, 5B**). This shared communication network likely reflects a universal tissue remodeling and wound healing response following the intense selective pressure of chemoimmunotherapy.

**Figure 5.**
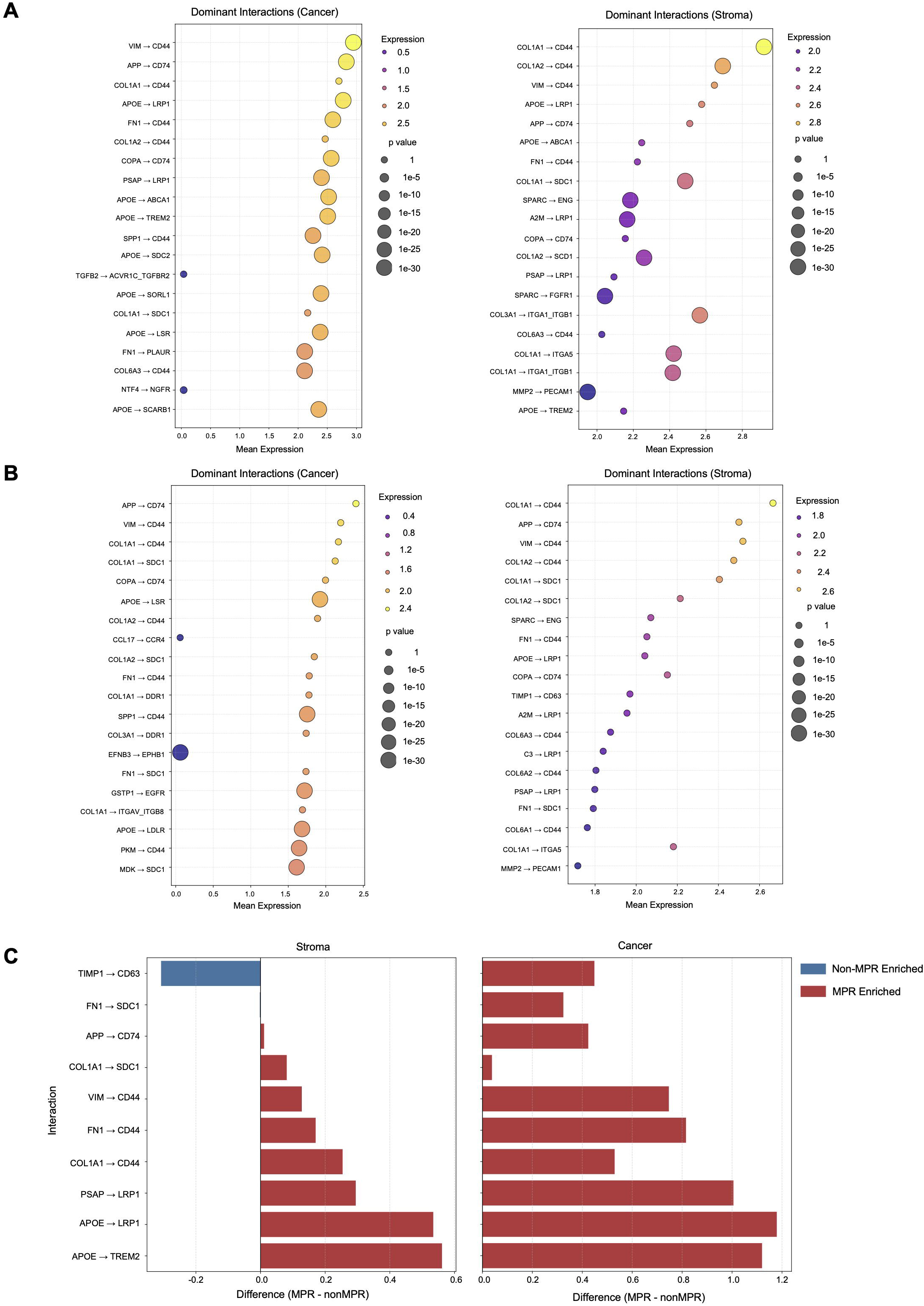
Spatially resolved ligand-receptor interactions in MPR and non-MPR tissues. (A-B) Dot plots show dominant ligand-receptor interactions in cancer and stroma regions from MPR (A) and non-MPR (B) tumors. The x-axis indicates mean expression of the ligand-receptor pair, and dot size reflects interaction significance. (C) Diverging bar plots compare selected interactions between groups in stroma and cancer regions; positive values indicate relative enrichment in MPR and negative values indicate relative enrichment in non-MPR. Matrix-associated CD44 interactions are observed in both response states, whereas interactions related to macrophage remodeling and scavenger pathways are relatively enriched in MPR, and TIMP1-CD63 is more prominent in non-MPR stroma.

However, the groups diverged significantly in their niche-specific communications. In the MPR tissues, the interaction network was uniquely enriched for macrophage-mediated clearance of apoptotic cells, known as efferocytosis. Specifically, signaling axes such as APOE–TREM2, PSAP–LRP1, and APP–CD74 were top-ranked in the MPR cancer regions (**Figure 5C**). The presence of these interactions suggests a microenvironment where macrophages actively clear dead cancer cells, thereby facilitating tissue resolution and deep immune infiltration.

In contrast, the non-MPR tissues exhibited a communication profile dominated by fibrotic barrier maintenance and cancer-intrinsic survival. Within the non-MPR stroma region, the TIMP1–CD63 axis was highly enriched. This interaction, coupled with extensive collagen signaling (e.g., COL1A1–SDC1, COL1A2–CD44), reinforces extracellular matrix remodeling and sustains the physical barrier that blocks effector T-cell penetration. Meanwhile, within the shielded non-MPR cancer core, interactions such as MDK–SDC1 and GSTP1–EGFR emerged as dominant signals. These axes represent an active resistance module, driving tumor cell survival, proliferation, and drug detoxification independent of immune regulation (**Figure 5B, 5C**).

Finally, we validated the structural organization of these communication networks using global spatial autocorrelation analysis (Moran’s I). The top L-R interaction scores for both the efferocytosis programs in MPR and the barrier/resistance modules in non-MPR demonstrated highly significant spatial clustering (Supplementary Figure 7). This confirms that these distinct communication pathways are not randomly distributed, but are rigidly structured into active functional niches, proving that the stromal barrier and resistance mechanisms are highly organized multicellular efforts.

### Co-localization of TROP2 and intrinsic resistance axes provides a spatial rationale for combination ADC therapies

Given the dense, immune-excluding stromal barrier and intrinsic chemo-resistance observed in non-MPR tumors, finding alternative therapeutic modalities is critical. ADCs targeting TROP2, an emerging therapeutic target in NSCLC, offer a rational strategy to physically bypass the stroma and deliver cytotoxic payloads directly to the epithelial compartment. To evaluate the spatial feasibility of this approach for treatment-refractory clones, we mapped TROP2 expression across the post-nCIT microenvironment.

Spatial expression analysis revealed that TROP2 was predominantly localized to the cancer regions, with minimal target expression in the surrounding stroma (**Figure 6A, 6B**). Notably, the expression level of TROP2 was significantly higher and more broadly distributed throughout the non-MPR resistant cancer cores compared to the residual cancer cells in the MPR group (**Figure 6A**). This spatial distribution confirms that TROP2 is structurally retained on these evolved resistant clones, identifying it as a potentially highly specific spatial anchor for direct drug delivery in non-responding patients.

**Figure 6.**
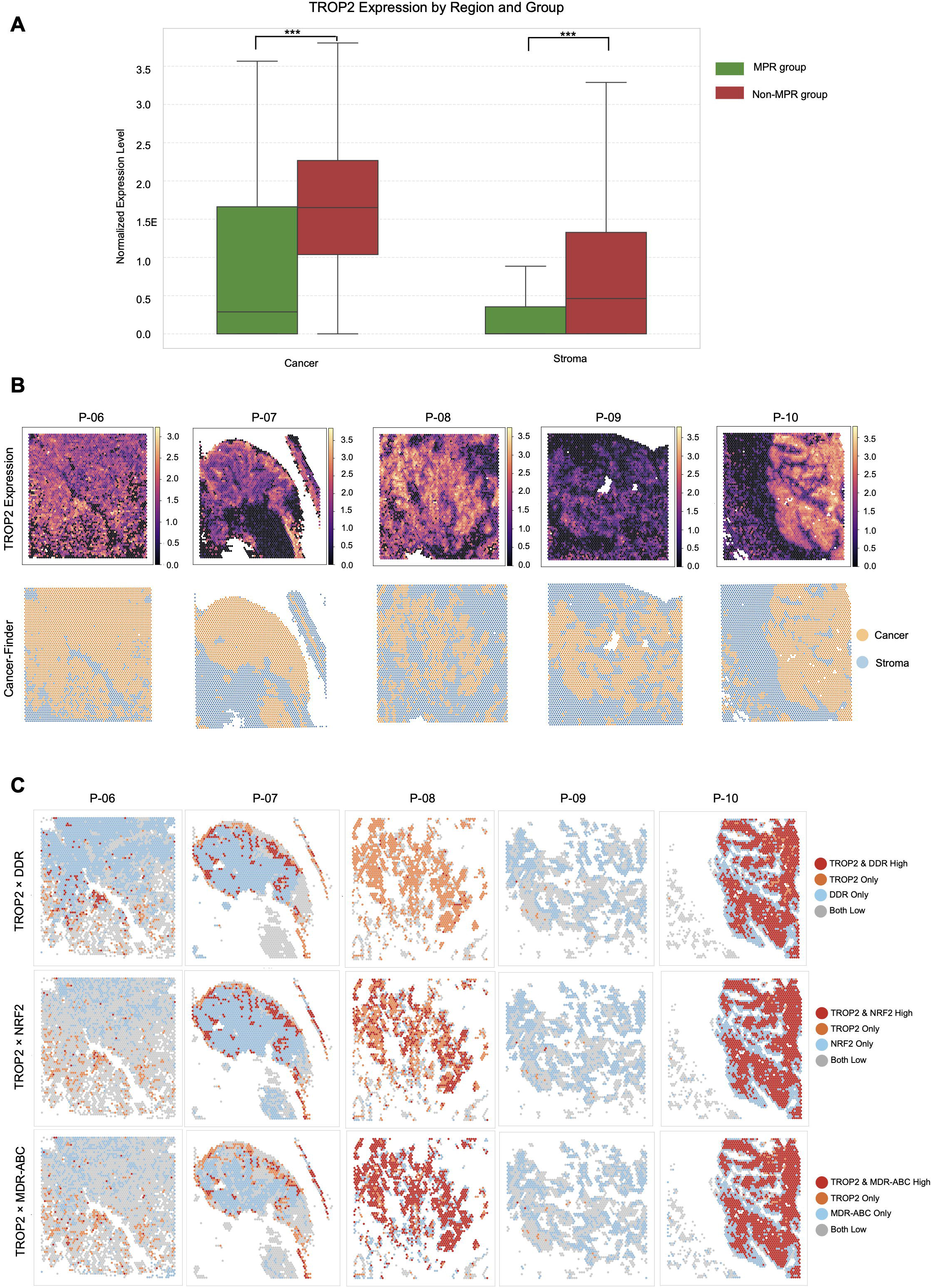
Spatial localization of TROP2 and its overlap with resistance-related programs. (A) Box plots compare TROP2 expression between MPR and non-MPR groups in Cancer-Finder-defined cancer and stroma regions. TROP2 expression was significantly enriched in the cancer regions of non-MPR tumors compared to both their corresponding stroma regions and the cancer regions of MPR tumors. Statistical significance was determined using the two-sided Wilcoxon rank-sum test (*** P < 0.001). (B) Representative non-MPR spatial maps show TROP2 expression together with region annotation. (C) Categorical co-localization maps display the spatial overlap of TROP2 with DDR-, NRF2-, and MDR-ABC-related signatures across non-MPR samples. These analyses show that TROP2 is retained in residual cancer regions and overlaps spatially with intrinsic resistance-related programs in non-MPR tumors, supporting its relevance as a candidate epithelial anchor for future residual disease-directed therapeutic evaluation.

However, successful payload delivery via ADCs must overcome the intrinsic survival programs previously identified within the cancer core. We therefore performed a spatial co-localization analysis between TROP2 expression and established resistance signatures. The mapping demonstrated a high degree of spatial overlap; regions strongly expressing TROP2 predominantly co-localized with high scores for DDR, NRF2-mediated antioxidant buffering, and MDR-ABC transporters (**Figure 6C**). These co-expression patterns indicate that while TROP2-ADCs can effectively guide therapeutic agents to the resistant epithelial core, the delivered payload will inevitably encounter a microenvironment primed with proficient DNA repair and detoxification mechanisms. This spatial evidence highlights the molecular landscape that the ADCs must navigate, providing a rationale for evaluating combination strategies to overcome the intrinsic repair capacity of the resistant niche.

## Discussion

Failure to achieve major pathologic response after neoadjuvant chemoimmunotherapy should not be interpreted simply as incomplete tumor regression. In resectable NSCLC, neoadjuvant chemoimmunotherapy improves clinically meaningful outcomes, and pathologic response is strongly linked to long-term survival; accordingly, the biology of residual non-MPR disease is not merely descriptive but clinically consequential [2,3,15]. Our data support a model in which non-MPR residual disease represents a spatially organized resistant niche selected under combined cytotoxic and immune pressure, rather than a passive remnant left behind by treatment. In this model, resistance is distributed across two cooperating compartments: a stromal boundary that constrains immune access and an epithelial core that remains optimized for survival.

One major implication of this framework is that persistent disease in non-MPR tumors appears to be driven by spatial immune exclusion rather than by immune scarcity alone. Prior spatial and single-cell studies in lung cancer have shown that distinct CAF states, matrix programs, and desmoplastic stromal architectures can marginalize T cells and reinforce immune suppression [16,17,21]. This concept is particularly relevant to a fibroblast-rich peripheral barrier and Treg-enriched interface in non-MPR disease. In lung adenocarcinoma, TIMP1 derived from tumor-associated fibroblasts can directly signal through CD63 on tumor cells to promote progression [18], and recent perioperative studies further suggest that fibroblast programs can impair neoadjuvant chemoimmunotherapy efficacy while fostering regulatory T-cell accumulation [19]. The boundary configuration in our cohort also fits with emerging evidence that CAFs and Tregs can operate as a cooperative immunosuppressive unit at the tumor interface [20]. By contrast, the responder state is better conceptualized as a remodeled tumor bed in which immune access and local antigen presentation are preserved or restored, rather than being spatially trapped outside the epithelial compartment [22].

Within that shielded core, the surviving epithelial compartment appears to undergo a functional shift away from immune engagement and toward autonomous endurance. Once immune access is spatially constrained, selection may favor cells that can repair therapy-induced DNA damage, neutralize oxidative stress, sustain mitochondrial metabolism, and maintain drug-detoxifying programs. Recent multi-omic work in neoadjuvant NSCLC has linked resistance to fibroblast programs and metabolic reprogramming [23,24], while experimental studies in lung cancer show that NRF2/GSH buffering, mtROS-linked plasticity, enhanced oxidative phosphorylation, and CYP-related detoxification can each contribute to chemoresistance [25–28]. In this context, the non-MPR core may represent a functionally decoupled compartment in which immune visibility becomes less relevant than preserving proliferative fitness under sustained therapeutic pressure. This interpretation moves the biology of non-MPR beyond simple pathway enrichment and toward a coordinated division of labor in which the stromal boundary excludes and the cancer core endures.

This spatial organization also has direct translational implications. If the dominant lesion in non-MPR disease is physical and functional separation from effector immunity, simply intensifying checkpoint blockade may have diminishing returns. A more rational strategy may be to bypass the stromal barrier and directly target the cancer core that remains structurally preserved after treatment. TROP2 is attractive in this setting because TROP2-directed antibody-drug conjugates have already shown clinical activity in previously treated NSCLC [29,30], and preclinical work demonstrates that TROP2-targeted TOP1 inhibitor ADCs can efficiently deliver DNA damage in TROP2-expressing tumors, including NSCLC models [31]. At the same time, the cells most accessible through a surface target may also be the cells best equipped to repair payload-induced injury. This creates a strong rationale for residual-disease-directed combinations that pair epithelial delivery with inhibition of DNA repair or related survival circuitry. Notably, preclinical studies and translational evidence from other solid tumors, including breast cancer, support the potential synergy between TROP2-directed TOP1 ADC strategies and DNA repair inhibitors such as PARP inhibitors [32,33]. While these combined approaches require further validation specifically in the context of residual NSCLC after nCIT, they provide a biological rationale for overcoming the intrinsic repair proficiency of the resistant niche.

Several limitations should be acknowledged. First, this study is transcriptomic and therefore cannot directly establish protein abundance, pathway flux, or metabolite activity. Second, although spatial transcriptomics preserves tissue architecture, the spot-based platform does not provide true single-cell resolution, and some niche-level interactions remain inferred rather than directly visualized at cell-cell contact level. Third, because our analysis is cross-sectional and restricted to post-resection specimens, it captures the clinically relevant endpoint of treatment persistence but not the full longitudinal evolution from pretreatment state to post-nCIT resistance. Fourth, the cohort is modest and histologically heterogeneous, which limits subgroup-specific inference. Finally, the proposed TROP2-based combination strategy remains hypothesis-generating and will require orthogonal protein-level validation and functional testing in matched ex vivo or in vivo models. In summary, our findings suggest that nCIT-refractory NSCLC is organized as a dual-compartment resistant niche composed of an immune-excluding stromal boundary and an epithelial core optimized for repair, detoxification, and continued survival. This view moves non-MPR beyond a response category and toward a biologically defined state of residual disease. More broadly, it provides a mechanistic framework for post-neoadjuvant precision strategies in resectable NSCLC—not only by targeting the surviving cancer core, but by dismantling the protective niche that allows it to persist.

## Declarations

### Ethics approval and consent to participate

This study was approved by the Institutional Review Board of Seoul National University Hospital (IRB approval number: H-2009-081-1158, approval date: Sep 18th, 2020) and conducted in accordance with the Declaration of Helsinki. Written informed consent was obtained from all participants prior to their inclusion in the study.

### Consent for publication

Not applicable

### Availability of data and material

The datasets used and/or analyzed during the current study are available from the corresponding author on reasonable request.

### Authors’ contributions

KJN, YTK, SHP, JK, and HC conceived and designed the study. SHP and KJN had full access to all the data in the study and take responsibility for the integrity of the data and the accuracy of the data analysis. JK performed the pathological assessment and provided annotations. HC and SB were responsible for the review of the research pipeline and technical validation. KJN, TY, JHP, BN, SAP, HJL, IKP, CHK, and YTK participated in patient data collection. SHP and KJN drafted the original manuscript. All authors participated in the acquisition, analysis, or interpretation of data, critically revised the manuscript for important intellectual content, and approved the final version of the manuscript. KJN and YTK supervised the study.

## Acknowledgements

We thank all members of Portrai, Inc. for the discussion and technical support.

## List of Abbreviations

ADC: Antibody-drug conjugate
APC: Antigen-presenting cell
DDR: DNA damage response
ECM: Extracellular matrix
EMT: Epithelial-mesenchymal transition
FAO: Fatty acid oxidation
FFPE: Formalin-fixed paraffin-embedded
GEE: Generalized estimating equations
GSH: Glutathione
HR: Homologous recombination
ICI: Immune checkpoint inhibitor
LISA: Local indicators of spatial association
MDR: Multi-drug resistance
MPR: Major pathologic response
nCIT: Neoadjuvant chemoimmunotherapy
NHEJ: Non-homologous end joining
NRF2: Nuclear factor erythroid 2-related factor 2
NSCLC: Non-small cell lung cancer
OXPHOS: Oxidative phosphorylation
ST: Spatial transcriptomics
TCA cycle: Tricarboxylic acid cycle
TME: Tumor microenvironment
Treg: Regulatory T cell

