## Supplementary files for "Distinct Spatial Programs of Response versus Resistance in Non-Small Cell Lung Cancer after Neoadjuvant Chemoimmunotherapy"

Supplementary information

Supplementary Figures

Supplementary Tables

**Supplementary Figure 1. Transcriptome-guided identification and spatial mapping of residual tumor regions.** (A) Representative H&E-stained images of post-nCIT NSCLC specimens illustrate the morphologic heterogeneity of treated tissues. (B) Spatial maps of viable residual tumor spots (indicated in orange) identified by the Cancer-Finder algorithm across MPR and non-MPR specimens, showing transcriptome-guided delineation of residual cancer regions from surrounding treatment-associated non-cancer tissue.

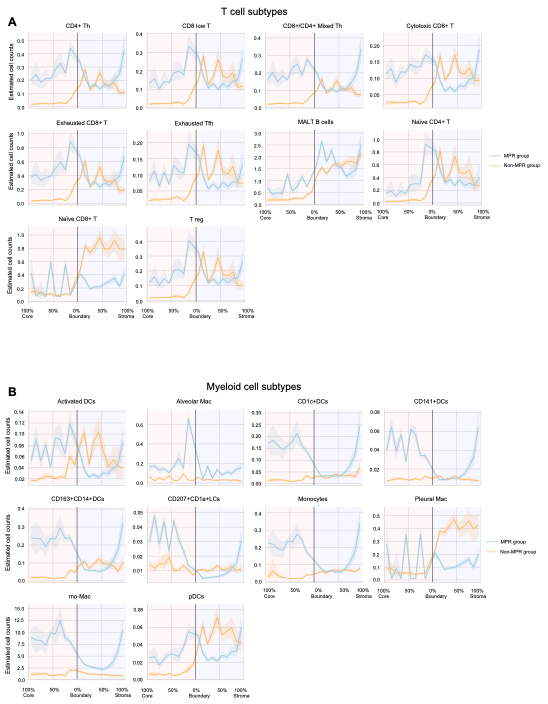

**Supplementary Figure 2. Detailed spatial distribution of T-cell and myeloid subtypes.** (A) Distance-based abundance profiles of T-cell subtypes estimated by cell2location along the normalized cancer-core-to-stroma axis, with the tumor boundary indicated by the central vertical line. Cytotoxic and exhausted CD8+ T-cell populations extend further toward the cancer-core side in MPR, whereas in non-MPR they are relatively enriched near the boundary and stromal side. (B) Corresponding profiles of myeloid subtypes. Activated dendritic cell and monocyte-related populations are more prominent on the cancer-core side in MPR, while these populations are comparatively reduced within non-MPR cancer regions.

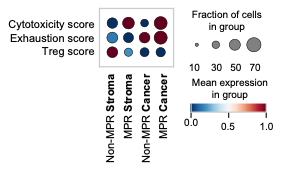

**Supplementary Figure 3. Regional distribution of immune functional scores.** Dot plots compare cytotoxicity, exhaustion, and regulatory T-cell (Treg) scores across MPR and non-MPR stromal and cancer compartments. Dot size indicates the fraction of spots within each group-region compartment, and color indicates the mean signature score. These patterns show relatively higher cytotoxicity and exhaustion scores in MPR compartments, particularly the cancer region, and relatively higher Treg scores in non-MPR stromal compartments.

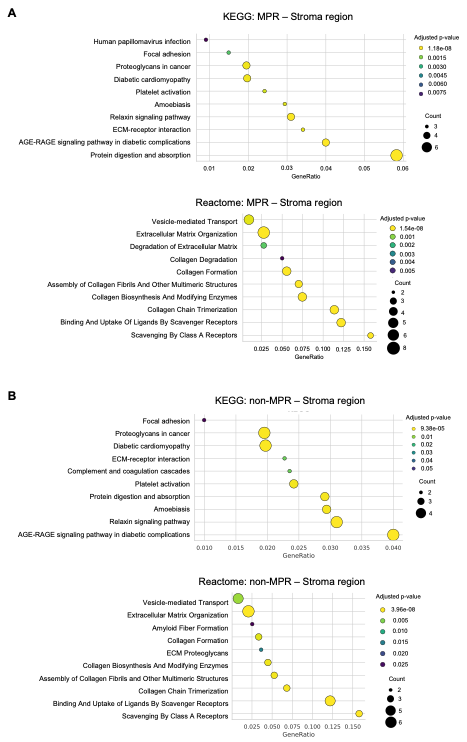

**Supplementary Figure 4. Pathway enrichment in stromal regions.** (A-B) KEGG and Reactome enrichment analyses for stromal regions from MPR (A) and non-MPR (B) specimens. The x-axis shows GeneRatio, dot size indicates gene count, and color indicates adjusted p value. Both groups show extracellular matrix- and collagen-related programs, whereas non-MPR stromal regions show greater representation of fibrotic and inflammatory pathways, consistent with a more matrix-rich stromal state.

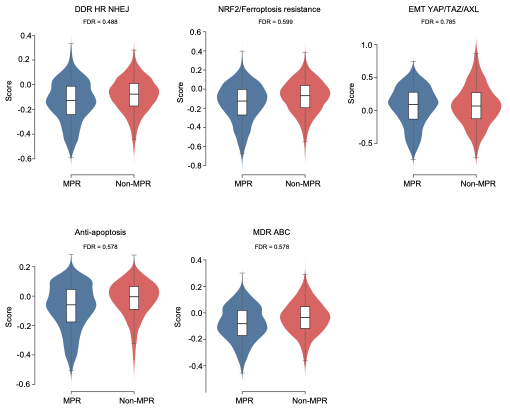

**Supplementary Figure 5**. Attenuated resistance-related signaling in stromal regions. Violin plots compare stromal signature scores between MPR and non-MPR groups for DNA damage response (DDR-HR/NHEJ), NRF2/Ferroptosis resistance, EMT YAP/TAZ/AXL, anti-apoptosis, and MDR-ABC programs. FDR values are shown above each comparison. Compared with the corresponding cancer-region analyses in the main figure, between-group differences are attenuated in the stroma, supporting enrichment of these programs within residual cancer regions rather than diffuse tissue-wide activation.

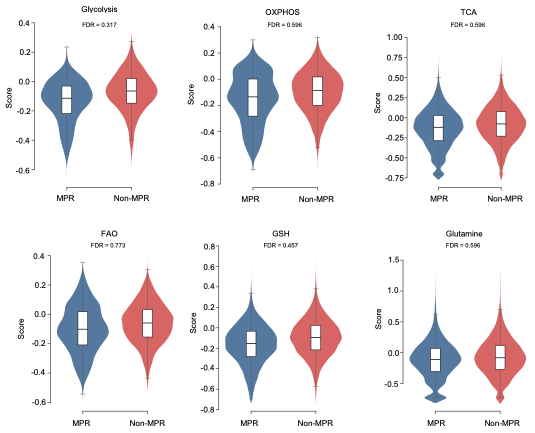

**Supplementary Figure 6. Attenuated metabolic rewiring in stromal regions.** Violin plots compare stromal pathway scores between MPR and non-MPR groups for glycolysis, oxidative phosphorylation, tricarboxylic acid cycle, fatty acid oxidation, glutathione metabolism, and glutamine metabolism. FDR values are shown above each comparison. Compared with the corresponding cancer-region analyses in the main figure, metabolic differences are attenuated in the stroma, suggesting that metabolic rewiring is concentrated in residual cancer regions.

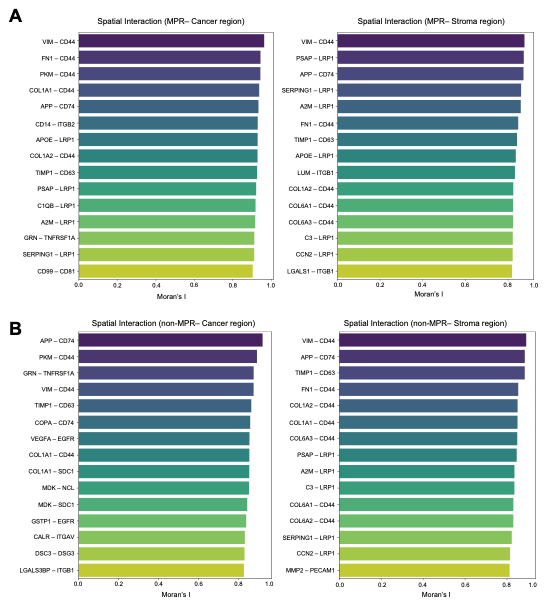

**Supplementary Figure 7**. Top spatially structured ligand-receptor interactions identified by Moran’s I. (A-B) Bar plots show the highest-ranked ligand-receptor pairs with significant spatial autocorrelation in MPR (A) and non-MPR (B) specimens, stratified by cancer and stromal compartments. Higher Moran’s I values indicate stronger spatial structuring of the interaction signal. The ranked interactions highlight distinct spatial communication landscapes across response states.

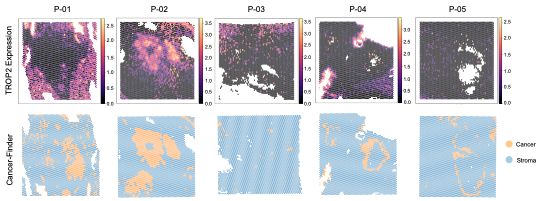

**Supplementary Figure 8. Spatial distribution of TROP2 in MPR specimens.** Top, spatial heatmaps of TROP2 expression across MPR tissues. Bottom, corresponding region maps showing Cancer-Finder-defined cancer and stroma regions. TROP2 signal is concentrated within cancer regions in MPR specimens, indicating that cancer-associated localization of TROP2 is maintained across response states.

**Supplementary Table 1.** **Differentially expressed genes (DEGs) stratified by spatial region.** The table provides a comprehensive list of the top DEGs identified when comparing MPR and non-MPR groups. To characterize niche-specific molecular profiles, the differential expression analysis was performed independently within the transcriptome-annotated cancer region and stroma region.

**Supplementary Table 2. Statistical details of enriched pathways.** This table provides the comprehensive statistical output, including raw p-values, false discovery rates (FDR), and Gene Ratios, for the Reactome and KEGG pathway enrichment analyses presented in the main figures. These metrics provide the statistical details underlying the Reactome and KEGG pathway enrichment analyses shown in the main figures.
